# Endurance Exercise Ameliorates Aging-Related Bradyarrhythmia in *Drosophila* Resulting from miR-283 Knockdown in LN_v_s

**DOI:** 10.1101/2025.02.25.640161

**Authors:** Qiufang Li, Xu Ping, Zhengwen Yu, Qin Yi, Chao Tang, Xiaoya Wang, Lan Zheng

**Affiliations:** Key Laboratory of Physical Fitness and Exercise Rehabilitation of Hunan Province, Hunan Normal University, Changsha, 410012, Hunan Province, China

**Keywords:** miRNA, aging, bradyarrhythmia, LN_v_s, exercise

## Abstract

MicroRNAs (miRNAs) are crucial in regulating cardiac aging and related diseases, yet few functional miRNAs have been identified. Prior studies showed miR-216a upregulation in heart failure patients, but its impact on aging hearts is unknown. Our study revealed systemic miR-283 overexpression or knockdown caused age-related bradycardia, mimicking human bradyarrhythmia. Importantly, we found that knockdown of miR-283 in ventral-lateral neurons (LN_v_s), rather than in the heart, led to the occurrence of bradyarrhythmia, which was mainly caused by the upregulation of miR-283 expression in the whole brain and heart. The gene of *clockwork orange* (*cwo*) may mediate miR-283’s effect on heart rhythm. Additionally, to investigate the miRNA regulatory mechanism underlying exercise-induced delay in cardiac aging, we conducted a three-week endurance exercise program on miR-283 knockdown flies in LN_v_s. We found that exercise significantly downregulated the accumulation of miR-283 in the brain and myocardium caused by aging or miR-283 knockdown in LN_v_s, improved the structure of myocardial fibers, and effectively reduced bradyarrhythmia. Our findings provides a new perspective on distal neuromodulation and intervention in cardiac aging.

## 1. Introduction

Normal aging hearts experience left ventricular hypertrophy, fibroblast proliferation, altered diastolic function, and decreased systolic reversal capacity, ultimately leading to decreased cardiac output and increased myocardial fibrosis [1]. This age-dependent change in cardiac structure and function is the main cause of the high prevalence of cardiovascular disease in the elderly population [1, 2], and is also the combined result of aging cardiovascular system dysfunction and autonomic nervous system imbalance [3–5]. Studies have shown that the imbalance of the renin-angiotensin system and sympathetic-vagal system homeostasis with age is an important trigger of cardiovascular aging [3, 5, 6], with the ultimate result being a decrease in the heart rate and myocardial contractility, increasing the risk of heart failure [3, 7]. Furthermore, research further reveals that changes at the cellular level are the main determinants of structural remodeling and functional changes in the aging heart [8–10]. MicroRNAs (miRNAs), a class of small non-coding RNAs, have been implicated at the post-transcriptional level by targeting hundreds of transcripts to regulate a variety of important cellular functions including stem cell self-renewal, cell proliferation, apoptosis, metabolism, and oxidative stress, and are considered to be regulators of cellular senescence [11, 12]. Several studies have shown that miRNAs play important roles in cardiovascular tissue homeostasis and cardiac aging processes. For example, age-dependent alterations of miR-34a [13], miR-195 [14, 15], and miR-217 [16] induce myocardial fibrosis, promote apoptosis, and cardiovascular dysfunction, and have been identified as circulating markers of de novo cardiac aging and cardiovascular risk, providing new perspectives for studying the biological processes of cardiac aging and therapeutic targets for delaying aging. We noted that circulating miR-216a is significantly upregulated in patients with heart failure [17] and induces endothelial senescence [18]. However, whether age-related decline in cardiac function is associated with altered miR-216a expression requires further investigation.

Aging and its consequences cannot be completely stopped; however, cardiac aging can be slowed down by appropriate means. Previous research has shown that appropriate physical exercise can reduce the incidence of age-related heart disease [19]. Rodent studies have further revealed the widespread benefits of exercise training in preventing apoptosis in the aging heart [20], reversing or “re-sensitizing” aged hearts to adrenergic stimulation [19, 21], improving Ca^2+^ cycling[22], and activating the IGF-1/PI3K/Akt signaling pathway [19]. Therefore, regular physical exercise has been proposed as a healthy lifestyle to delay aging and prevent and treat diseases [23–25]. In recent years, an increasing number of studies have found that exercise protects heart health and prevents cardiac dysfunction by regulating miRNAs with apoptotic potential [26–30]. However, the role of miRNAs in exercise-improving function in aging hearts remains largely unknown.

The *Drosophila* heart, also referred to as the dorsal vessel, has some conserved regulatory mechanisms related to vertebrate heart development and aging [31], and has been widely used to study the physiological, genetic, and epigenetic basis of cardiac aging [32]. Furthermore, the detection of mature heart function in *Drosophila* allows the accurate assessment of the functional characteristics of aging hearts [33, 34]. Exercise based on the innate anti-gravity climbing trait of *Drosophila* has been shown to be prominently effective in preventing age-related cardiac dysfunction [35–38]. In this study, we used a *Drosophila melanogaster* model to reveal the regulatory role of dme-miR-283 (miR-283, a homolog of miR-216a in *Drosophila*) in age-dependent cardiac function decline, and further explored whether the ameliorative effects of exercise on the aging heart are associated with exercise resistance to age-dependent changes in miR-283 expression.

## 2. Results

### 2.1 Functional and structural deterioration of the myocardium in aging *Drosophila*

Analysis of cardiac function in aging *w^1118^* flies (10 days of age corresponding to the young stage, 30 days of age for middle age, and 50 days of age for old age) [39] using SOHA revealed that a significant reduction in HR and an increase in HP were already present at midlife and persisted into old age (Figure 1A and 1B), mainly caused by the prolonged DI and SI (Figure 1C and 1D). M-mode traces from the SOHA showed a more prevalent incidence of bradycardia in middle-aged and older hearts (Figure 1F and 1G). The occurrence of diastolic insufficiency was also significantly increased compared to that in younger flies (Figure 1I), suggesting a more abnormal cardiac rhythm in middle-aged and older *Drosophila*. In addition, another change with aging was manifested by narrowing of the heart lumen canal diameter, with significant shortening of SD and DD at 30d and 50d (Figure 1F and 1G), showing an increase in FS due to a greater narrowing of the SD (Figure 1J) but a significant decrease in cardiac output (Figure 1K). Moreover, the incidence of incomplete relaxations was also significantly increased in *Drosophila* at 30d and 50d (Figure 1F and 1G). Myocardial SA-β-gal staining also showed a significant increase in a blue precipitated area at 30d that persisted until 50d (Figure S 1A-1D). Therefore, our results show that *Drosophila* has already experienced comprehensive age-related deterioration of cardiac rhythm and pumping capacity at 30d, and this deterioration continues until 50d.

**Figure 1.**
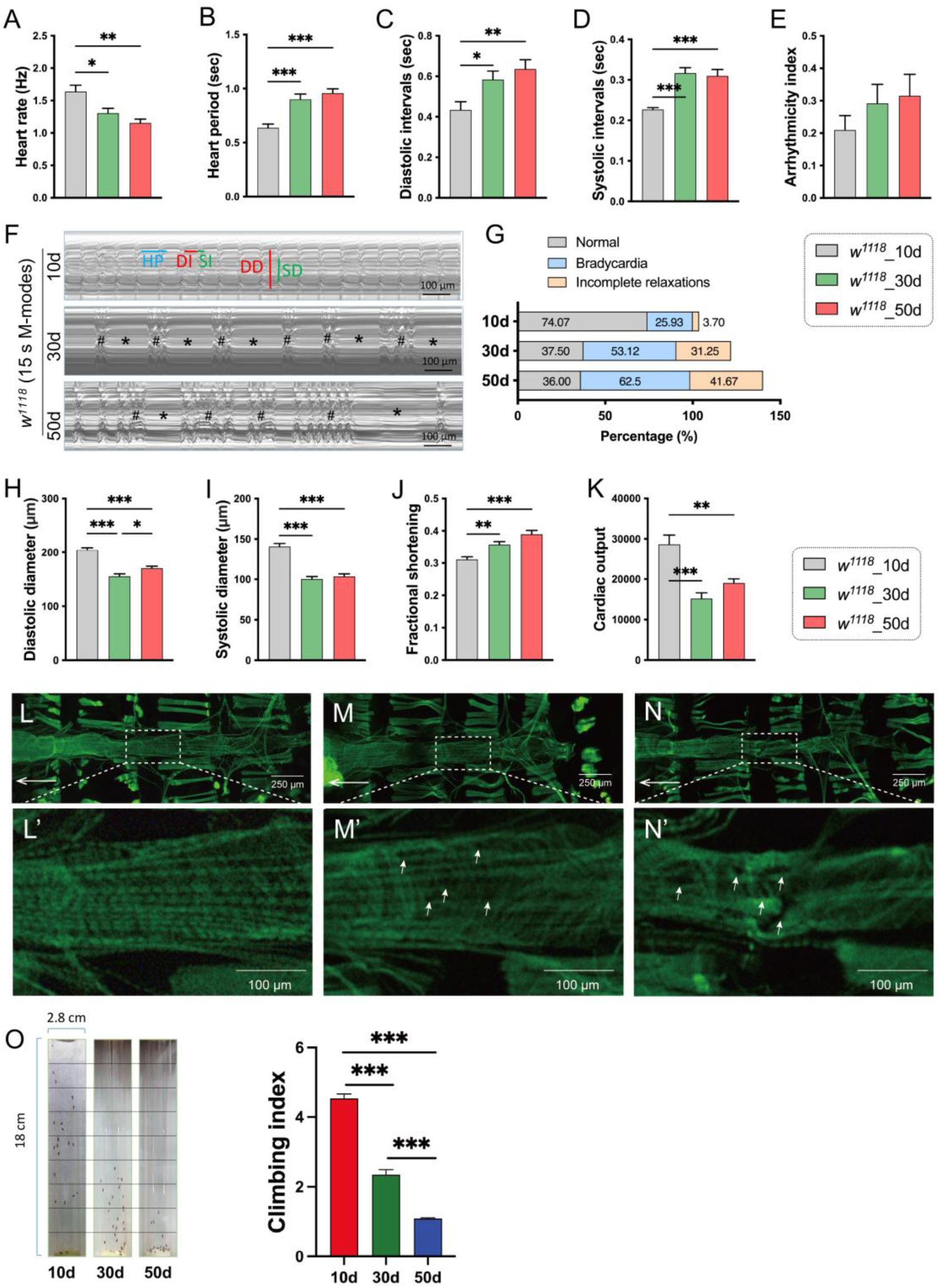
Changes in cardiac function and structure during the aging process in *Drosophila*. (A-E) Heart rate (HR), heart period (HP), diastolic interval (DI), systolic interval (SI), and arrhythmia index (AI) in *w^1118^* 10-day-old, 30-day-old, and 50-day-old *Drosophila*. (F) Representative M-mode traces of heartbeats in 10d, 30d, and 50d *Drosophila*. The blue horizontal line represents a complete cardiac cycle, with red indicating the DI and green indicating the SI. Red vertical lines represent diastolic diameter (DD), and green vertical lines represent systolic diameter (SD). “*” indicates bradycardia, and “#” indicates incomplete relaxations. Scale bar: 100 μm. (G) Percentage of *Drosophila* with a diastolic interval (DI) greater than 1 second (indicating bradycardia) and percentage with a DI less than 0.06 seconds (indicating incomplete relaxations), calculated based on M-mode traces. (H-K) DD, SD, fractional shortening (FS), and cardiac output (CO) in 10d, 30d, and 50d *Drosophila*. Cardiac function measurements: n=25-35. (L-N) Phalloidin-stained images of *Drosophila* hearts acquired using the Leica somatic fluorescence microscopy. The arrow points in the direction of the head of the fruit fly. Magnified images of the heart tube at the 2nd-3rd abdominal segments are shown in L’-N’, corresponding to the regions imaged by M-mode. Prominent myofibrillar gaps are indicated by white arrows. n=5-8. (O) Left: Instantaneous climbing height at the 5th second for 10d, 30d, and 50d *Drosophila*. The climbing tube is divided into nine equal sections, with scores ranging from 1 to 9 from bottom to top. The average climbing score for each group is calculated as the climbing index (right). n=100-150. One-way ANOVA followed by Tukey multiple comparisons test. *** p < 0.001, ** p < 0.01, * p < 0.05.

Visualization of cardiac filamentous actin (F-actin) by Phalloidin staining similarly showed an increase in myofibrillar gaps in middle-aged *Drosophila* hearts and a larger staining deficit in aged myocardium (Figure 1L-1N). These results suggest a progressive deterioration of *Drosophila* myocardial structure with age, which may contribute to the decline in myocardial pumping function and the increase in arrhythmias.

Meanwhile, we observed a significant enhancement of SA-β-gal activity in the brain at 30d that persisted until 50d (Figure S 1E-1H), a change similar to that of aging myocardium. And the climbing ability was continuously decreased with age at 30d and 50d (Figure 1O), and this age-related decline in behavior also suggests that *Drosophila* undergoes neurodegeneration during aging [40, 41]. Our study reveals that age-dependent deterioration of cardiac function with neurologic deficits is already occurring in *Drosophila* at the age of 30d.

### 2.2 Dysregulation of miR-283 expression triggers aging-related cardiac functional alterations in young *Drosophila*

To determine whether miR-283 regulates cardiac function, we examined cardiac performance in young miR-283 heterozygous knockout (mir-283^KO/+^), homozygous knockout (mir-283^KO^) and overexpressing (*Tub*>mir-283^OE^) flies. And whole-body miR-283 expression levels were examined (Figure 2A, whole body), while the myocardium and brain also showed consistent partial knockdown, complete knockout, and overexpression (Figure 2A, heart and brain). Analysis of heart rhythm indices found that both miR-283 knockout and overexpression resulted in decreased HR (Figure 1B), prolonged HP and DI (Figure 1C and 1D), and a general increase in bradycardia (Figure 1G and 1H), but did not induce an aging-associated increase in SI (Figure 1E). These results suggest that miR-283 knockout and overexpression lead to an increased incidence of long DI-associated bradycardia with reduced heart rate, which is phenotypically similar to bradyarrhythmia defined clinically on the basis of heartbeat rate[42, 43]. Moreover, differing from miR-283 knockout, miR-283 overexpression also resulted in an overarching reduction in pumping capacity (Figure 1I-1L), as well as a decrease in the climbing index (Figure 1M), suggesting that systemic overexpression of miR-283 produced more widespread negative effects. Additionally, it is worth stating that we drove miR-283 systemic overexpression with *Tub*-Gal4, which generated a particularly high lethality rate, further suggesting that systemic increase in miR-283 expression triggers severe consequences.

**Figure 2.**
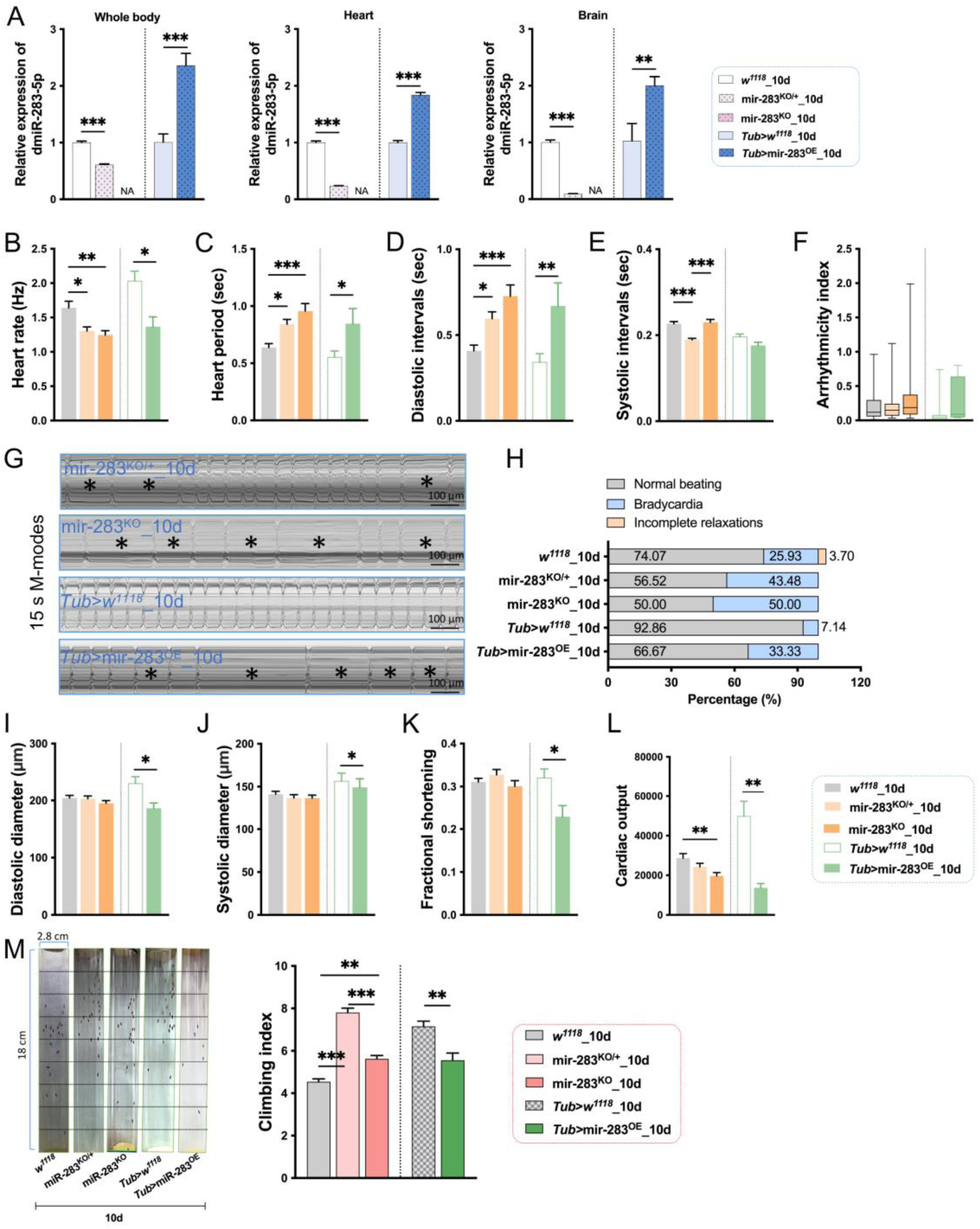
Effects of miR-283 knockout and overexpression on heart function and climbing ability. (A) RT-qPCR analysis of miR-283-5p expression levels in the whole body, heart muscle, and brain of control (*w^1118^*), miR-283 heterozygous knockout (mir-283^KO/+^), homozygous knockout (mir-283^KO^), as well as control (*Tub*> *w^1118^*) and miR-283 overexpression (*Tub*>mir-283^OE^) *Drosophila*. Whole body: n=3-5; heart muscle and brain: n=20-30. (B-H) Heart rhythm-related indicators. Scale bar: 100 μm. (I-L) Heart pumping capacity-related indicators. n=25-35. (M) Left: Representative climbing heights at the 5th second for each group of *Drosophila*. Right: Corresponding climbing index. n=100-150. Exceptionally, *Tub*>mir-283^OE^ group had a high lethality rate, and thus the number of flies available for testing was n=6. One-way ANOVA followed by Tukey multiple comparisons test was used for comparisons between *w^1118^*, mir-283^KO/+^, and mir-283^KO^; Student t-test was used for comparisons between *Tub*> *w^1118^* and *Tub*>mir-283^OE^. *** p < 0.001, ** p < 0.01, * p < 0.05.

We next explored whether the slowing of heart beat caused by altered miR-283 expression was consistent with an age-induced decrease in HR. We performed a two-way ANOVA for gene*age on HR, HP and DI in mir-283^KO/+^ and mir-283^KO^ senescent *Drosophila* and found a significant interaction between miR-283 knockout and aging (Table S2). In addition, there was a significant rise or trend towards an increase in HP and DI in 10d and 30d miR-283 knockout flies (Figure S2B-S2C), which resulted in a decrease in HR (Figure S2A). Collectively, these findings suggest that miR-283 regulates age-related HR reduction and DI prolongation, and that dysregulation of its expression may contribute to the development of bradycardic arrhythmias.

### 2.3 miR-283 knockdown in LN_v_s, but not heart, leads to bradyarrhythmia

We examined changes in miR-283 expression in senescent myocardium, including wild-type *w^1118^* and hybridized controls (*w^1118^*>mir-283^SP^). The results showed that the expression of miR-283 in the myocardium of *w^1118^* flies exhibited a dynamic change of decrease at 30 d and increase at 50 d (Figure S3A). In contrast, *w^1118^*>mir-283^SP^ *Drosophila* showed a dynamic change of increase at 30d and decrease at 50d (Figure S3B). This difference may be related to the fact that these two types of *Drosophila* are in different life states. Typically, wild-type *w^1118^* “aged” faster than the hybridized control. Overall, however, our results suggest that myocardial miR-283 expression changes during aging.

To determine whether age-related decline in cardiac function is associated with downregulation of myocardial miR-283 expression, we used the GAL4/UAS system to express mir-283 sponge in the heart, resulting in a specific knockdown of miR-283. We crossed UAS-mCherry.mir-283.sponge virgin strains with heart-specific tissue drivers (*Hand*-Gal4) and observed that the cardiomyocytes and paracardiocytes of the offspring generation were labeled with mCherry fluorescence (Figure 3A), and the expression of miR-283 in the myocardial tissue was significantly reduced (Figure 3N), indicating that myocardial successful knockdown of miR-283.

**Figure 3.**
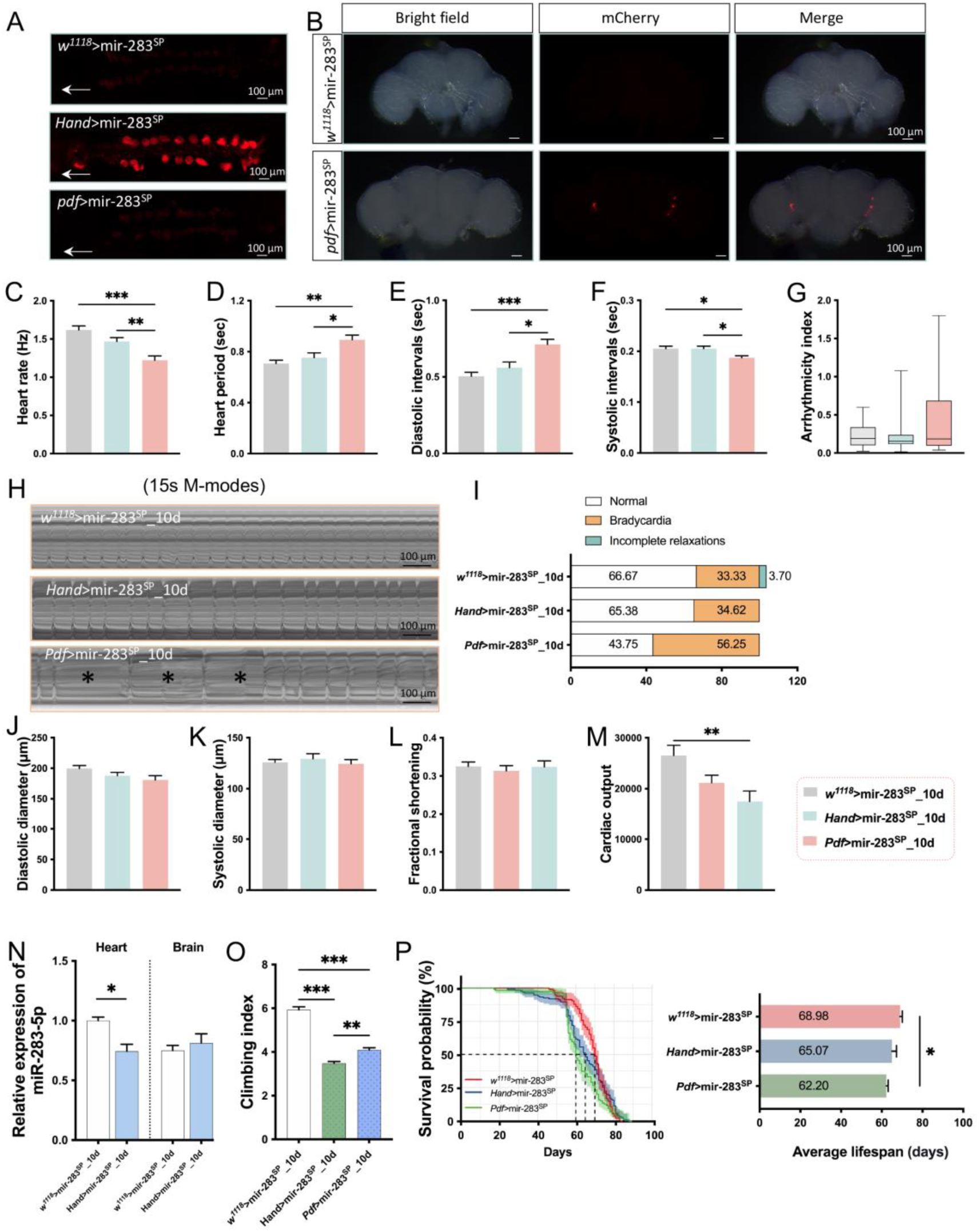
Effects of miR-283-specific knockdown in myocardium or LN_v_s on cardiac function, climbing ability, and lifespan. (A) Virgin flies expressing UAS-mCherry.mir-283.sponge were crossed with *w^1118^*, *Hand*-Gal4, and *Pdf*-Gal4 male flies. Spontaneous fluorescence images of the myocardium in the F1 generation were captured using a Leica stereomicroscope. White arrows points in the direction of the head of the flies. (B) Virgin flies expressing UAS-mCherry.mir-283.sponge were crossed with *w^1118^* and *Pdf*-Gal4 male flies. Brightfield (left), spontaneous fluorescence (middle), and merged (right) images of the *Drosophila* brain in the F1 generation were captured using a Leica stereomicroscope. Scale bar: 100 μm. (C-I) and (J-M) Represent heart rhythm-related indicators and heart pumping capacity-related indicators, respectively. n=20-35. One-way ANOVA followed by Tukey multiple comparisons test. *** p < 0.001, ** p < 0.01, * p < 0.05. Scale bar: 100 μm. (N) RT-qPCR analysis of the relative expression levels of miR-283-5p in the myocardium of miR-283 knockdown flies (*Hand*>mir-283^SP^) compared to the control group (*w^1118^*>mir-283^SP^) using U6 as the reference gene. n=20-30. Student t-test. * p < 0.05. (O) Climbing index of control, myocardial miR-283 knockdown, and LN_v_s miR-283 knockdown flies. n=110-200. One-way ANOVA followed by Tukey multiple comparisons test. *** p < 0.001, ** p < 0.01. (P) Survival curves (left) and mean lifespan (right) of control, myocardial miR-283 knockdown, and LN_v_s miR-283 knockdown flies. n=115-135. One-way ANOVA followed by Tukey multiple comparisons test. * p < 0.05.

Next, we examined cardiac function in young *Hand*>mir-283^SP^ *Drosophila*. Compared to age-matched controls (*w^1118^*>mir-283*^SP^*), myocardial tissue miR-283 knockdown did not have a significant effect on heart beating and contractility (Figure 3C-3M). This indicates that partial decrease of miR-283 in the heart does not affect cardiac function or that there are other compensatory regulatory mechanisms to maintain normal performance of the heart. mir-283^KO/+^ *Drosophila* exhibited features of bradyarrhythmia along with a significant reduction in miR-283 expression in the myocardium and brain (Figure 2A-2L). However, specific knockdown of myocardial miR-283 did not affect cardiac function (Figure 3C-3M). Therefore, we hypothesized that the cardiac phenotype of mir-283^KO/+^ *Drosophila* may be regulated by neuronal miR-283.

In addition to its own pacing regulation, the heartbeat is also regulated by the “brain-heart” axis from the vagus and sympathetic nerves of the brain [44]. The mechanism of brain-heart regulation is based on the hierarchical clock system, where the main biological clock that coordinates the circadian activity of the whole body is located in the suprachiasmatic nucleus (SCN) of the hypothalamus, which transmits neurotransmitters to the vagus and sympathetic nerves through an anatomical connection with the paraventricular nucleus of the hypothalamus. This enforces endogenous rhythm enforcement to the heart and other organs, and regulates HR and rhythm via the sinus node [44]. The *Drosophila* circadian pacemaker is located in LN_v_s of the brain. We used LN_v_s-expressing Gal4 (*Pdf*-Gal4) to drive the mir-283 sponge to investigate whether miR-283 regulates cardiac rhythm through the brain-heart axis. We observed mCherry fluorescence in the LN_v_s of *Pdf*>mir-283^SP^ *Drosophila* (Figure 3B), indicating successful driving. The results of cardiac function in young *Drosophila* with *Pdf*>mir-283^SP^ showed that 10d flies exhibited a significantly reduced HR (Figure 3C) and prolonged HP and DI (Figure 3D - 3E) heartbeat pattern, especially a significantly increased incidence of bradycardia (Figure 3H - 3I), which is consistent with the senescent heartbeat phenotype induced by mir-283 knockdout or overexpression. Similarly, we did not find a function of miR-283 in LN_v_s to regulate cardiac contraction (Figure 3J - 3L), but resulted in a significant reduction in cardiac output due to decreased HR (Figure 3M).

Furthermore, we observed a decline in climbing ability in *Hand*>mir-283^SP^ and *Pdf*>mir-283^SP^ 10d flies (Figure 3O), suggesting that imbalances in miR-283 homeostasis in the heart or brain accelerate neurosenescence. A leftward shift in the survival curve of *Hand*>mir-283^SP^ and *Pdf*>mir-283^SP^ *Drosophila* was also observed (Figure 3P), especially a significant decrease in the median survival time (Figure 3P) and mean lifespan (Figure 3P) in *Pdf*>mir-283^SP^ flies, further suggesting that stable expression of miR-283 in LN_v_s is important for healthy aging.

We further analyzed the effect of aging combined with miR-283 knockdown in LN_v_s on heartbeat rhythm in *Drosophila*. Results of a two-way ANOVA on aging and miR-283 knockdown showed an interaction between aging and knockdown in HR, HP, and DI (Table S3). Senescence changes with reduced HR, prolonged HP and DI consistent with wild type were observed in control group (*w^1118^*>mir-283^SP^, Figure S3). Meanwhile, *Pdf*>mir-283^SP^ exhibited cardiac senescence characteristics (significantly reduced HR and prolonged HP and DI) in youth and persisted into middle age (Figure S3). In conclusion, our findings suggest that specific knockdown of miR-283 in LN_v_s, but not in the myocardium, leads to an age-associated decrease in HR, along with an increased incidence of DI prolongation-associated bradycardia, and ultimately an increase in bradyarrhythmia.

### 2.4 Knockdown of miR-283 in LN_v_s induces heart and brain aging and impairs cardiac morphological

We next examined the effect of miR-283-specific knockdown in LN_v_s on miR-283 expression in the myocardium and brain. qPCR results showed a significant increase in miR-283 expression in the brain and myocardium of 10d *Pdf*>mir-283^SP^ *Drosophila*, which compared with age-matched controls (*w^1118^*>mir-283^SP^) (Figure 4A). To determine whether this increased expression was consistent with changes in aging, we further examined miR-283 expression levels in 10d and 30d *Drosophila*. The results revealed an elevation of miR-283 expression in both myocardium and brain of *Drosophila* in the 30d *w^1118^*>mir-283^SP^ and *Hand*>mir-283^SP^ groups, as compared to their 10d counterparts of the same genotype (Figure 4A). Brain miR-283 expression was also significantly increased in the senescent *Pdf*>mir-283^SP^ *Drosophila*, whereas a high level of expression consistent with 10d was maintained in the myocardium (Figure 4A). These results suggest that miR-283 knockdown in LN_v_s leads to an accumulation of miR-283 expression in the myocardium and brain consistent with senescence.

**Figure 4.**
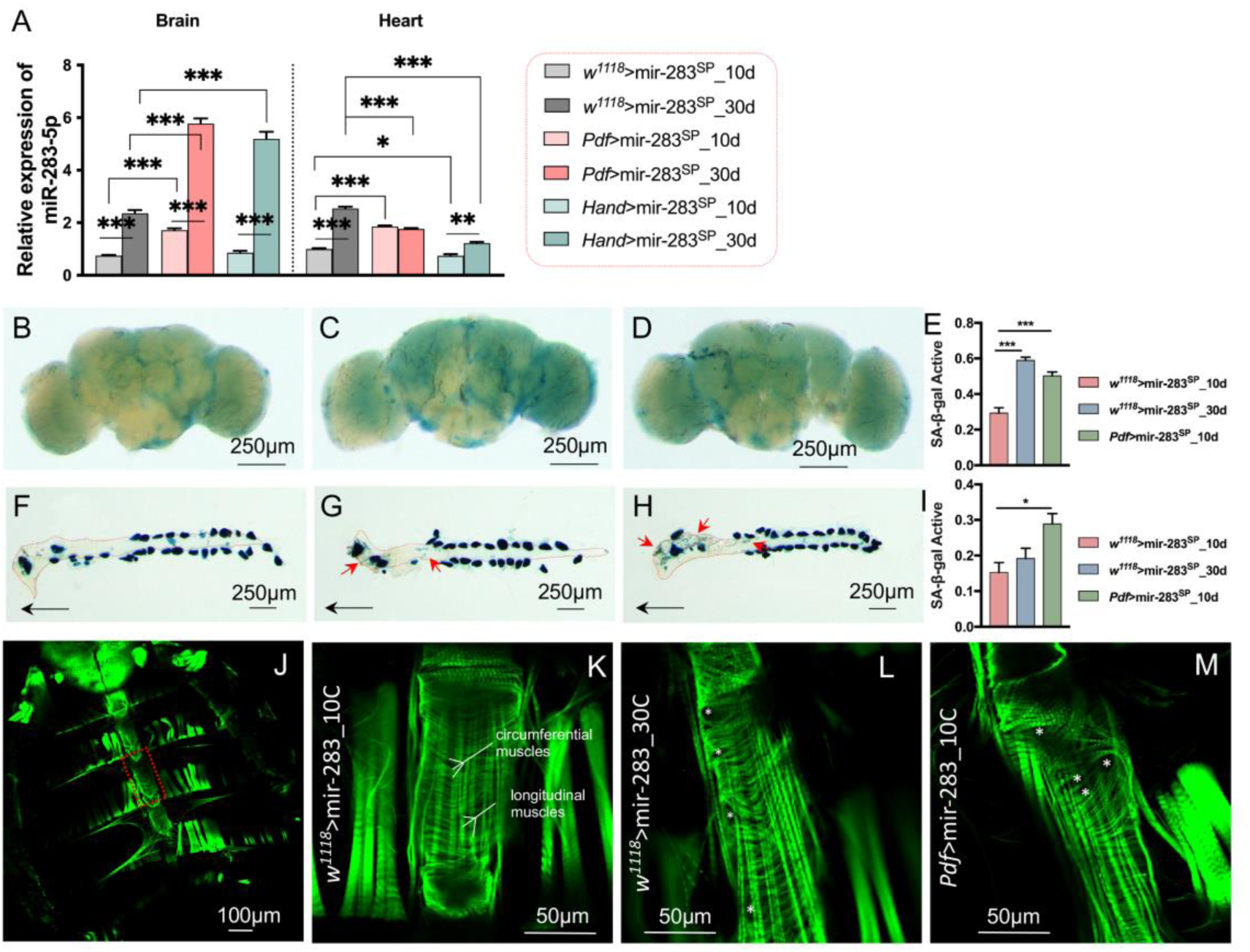
miR-283 Knockdown in LN_v_s triggers brain and heart aging. (A) RT-qPCR analysis of miR-283 expression levels in the myocardium and whole brain of 10d and 30d control (*w^1118^*>mir-283^SP^), myocardial miR-283 knockdown (*Hand*>mir-283^SP^), and LN_v_s miR-283 knockdown (*Pdf*>mir-283^SP^) *Drosophila*. n=20-30. (B-D) Representative images of SA-β-gal staining in the brain of *w^1118^*>mir-283^SP^ flies at 10d and 30d, and(*Pdf*>mir-283^SP^ flies at 10d. n=5-8. (E) Percentage of SA-β-gal-positive staining in the brain analyzed using Image J. (F-H) Representative images of SA-β-gal staining in the heart of *w^1118^*>mir-283^SP^ flies at 10d and 30d, and *Pdf*>mir-283^SP^ flies at 10d. The outline of the heart tube is outlined with a red dashed line. Black arrows points in the direction of the head of the flies. Red arrows point to significant SA-β-gal precipitate areas. n=5-8. (I) SA-β-gal-positive staining was analyzed as a percentage of the total myocardium using Image J. One-way ANOVA, * p < 0.05. (J) Complete myocardium Phalloidin staining and imaging. The red dashed box corresponds to the M-mode capture area, which was imaged at 600× magnification to obtain (K-M). Transverse circular fibers represent the muscles responsible for myocardial contractile function, while longitudinal fibers represent the ventral longitudinal muscles. Significant gaps between myofibrils are indicated with“*”. n=5-8. One-way ANOVA followed by Tukey multiple comparisons test. *** p < 0.001, ** p < 0.01.

Senescence-associated β-galactosidase (SA-β-gal) is a marker of aging cells in vitro, vertebrates, and *Drosophila*, and histochemical assays are considered the most reliable and easily detectable age-dependent markers [45, 46]. To further characterize that miR-283 knockdown in LN_v_s induces senescence, we stained brain and heart tissues of *w^1118^*>mir-283^SP^ (10d and 30d) and *Pdf*>mir-283^SP^ (10d) *Drosophila* using SA-β-gal. The results showed a significant increase in blue precipitated areas in 30d brains of *w^1118^*>mir-283^SP^ as well as in the 10d brains of *Pdf*>mir-283^SP^ (Figure 4B-4E), indicating that both aging and miR-283 knockdown in LN_v_s resulted in enhanced SA-β-gal activity throughout the brain. In senescent hearts, conical chambers and within the walls of the heart tube showed sporadic granular staining (Figure 4G), unlike in LN_v_s knockdown of miR-283-induced senescent hearts, where stained regions covered most of the conical chambers and heart tube (Figure 4F-4I), further suggesting that knockdown of miR-283 in LN_v_s simultaneously induces heart and brain aging.

The adult *Drosophila* heart consists of circumferential muscles, longitudinal muscles, and alary muscles, of which the circumferential muscles are the working cells of the heart [47]. To investigate whether aging and miR-283 knockdown in LN_v_s result in structural changes in filamentous actin (F-actin) in myogenic fibers, we stained semi-exposed hearts with phalloidin and obtained 600-fold magnified images of cardiac structures at the third ventral segment (Figure 4J). In control young hearts, myogenic fibers were circular and tightly aligned (Figure 4K) to promote consistent cardiac contraction. While in aged *w^1118^*>mir-283^SP^ *Drosophila* (Figure 4L) as well as in young *Pdf*>mir-283^SP^ *Drosophila* (Figure 4M), consistent disorganization of myogenic fiber arrangement and increased gaps were observed. This suggests that both aging and miR-283 knockdown in LN_v_s lead to structural degradation of myogenic fibers, which may contribute to the development of bradyarrhythmias.

### 2.5 Bioinformatics prediction and experimental validation of potential targets of miR-283

*Drosophila* miR-283 is located on the intron of the X-chromosome-encoded gene Gmap, and the pre-mir-283 is processed to generate the mature miR-283-5p, which is conserved with miR-216a in species such as humans, rats, and mice (Figure 5A). Previous studies have shown that miR-216a may play a pivotal role in aging cardiovascular diseases (atherosclerosis and heart failure) [18] and has been proposed as a new biomarker for the diagnosis of heart failure and related diseases [17], but its regulatory mechanisms remain unclear. We predicted by DIANA microT-CDS online tool that miR-283-5p target genes are mainly enriched in Notch signaling pathway (*Nct*, *mam*, *Ser*, *Delta*, *Hairless*) (Figure 5B). In our study, the expression level of miR-283 in the brain and heart of 10d *Pdf*>mir-283^SP^ flies exhibited a consistent increase with that of the control aging *Drosophila* (Figure 5C). Examination of the predicted target genes of the Notch signaling pathway showed that *Nct*, *Ser* and *mam* were significantly downregulated in both senescent and *Pdf*>mir-283^SP^ brains. However, the predicted target genes were significantly upregulated or tended to increase in both aging and *Pdf*>mir-283^SP^ hearts (Figure 5D-H) and were not consistent with the expected effect of miRNA sponge inhibition.

**Figure 5.**
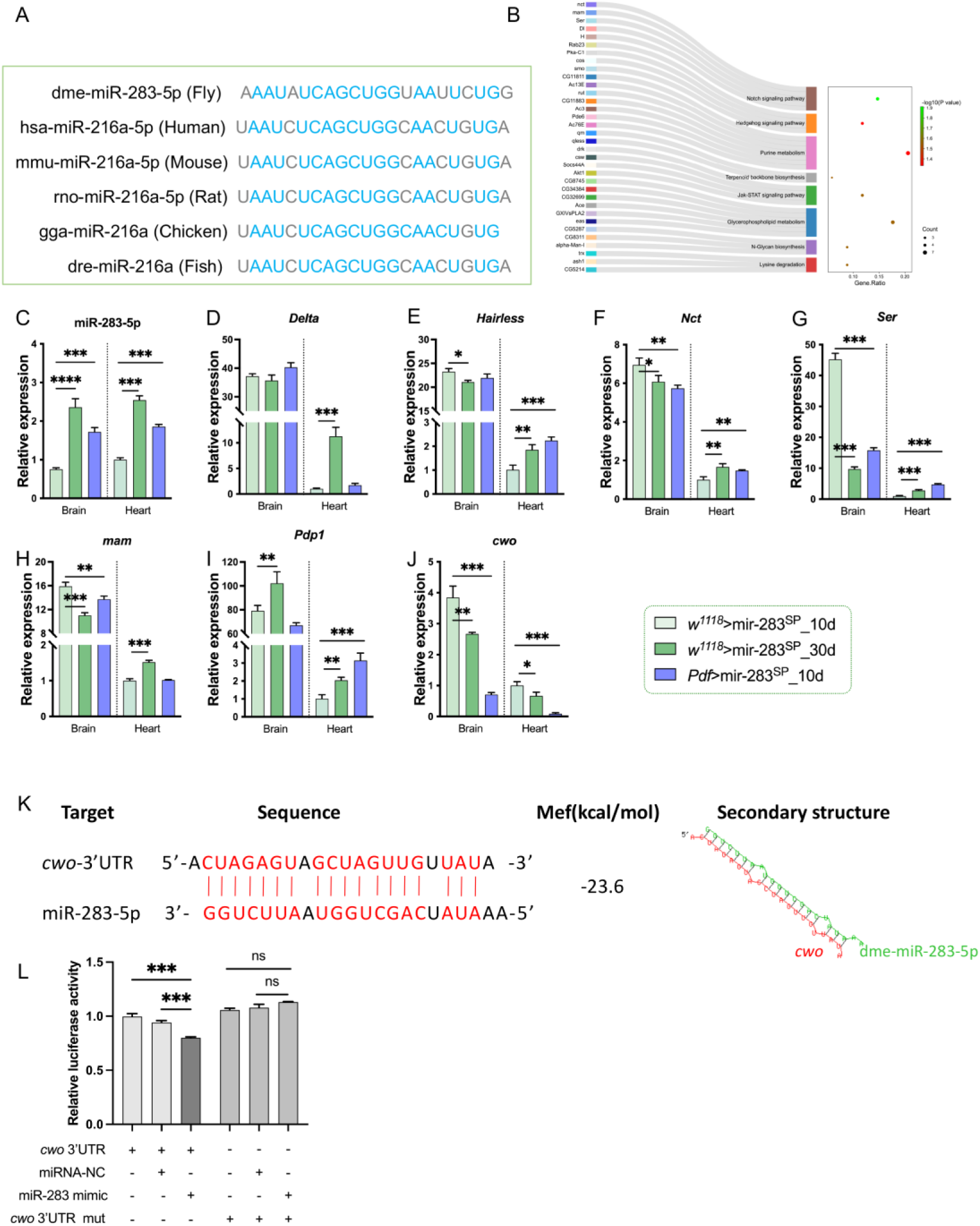
Prediction and validation of potential target genes of miR-283. (A) Homology alignment of the *Drosophila* miR-283-5p sequence with other species. (B) Prediction and enrichment analysis of target genes of miR-283-5p using the DIANA microT CDS online tool. (C-J) qPCR analysis of the effects of aging (10d and 30d *w^1118^*>mir-283^SP^) and miR-283 knockdown in LN_v_s (*Pdf*>mir-283^SP^) on the expression levels of miR-283-5p and its predicted target genes in the brain and heart. n=20-30. (K) Prediction of the targeted binding site between miR-283-5p and the 3’UTR of *cwo* using the RNAHybrid online tool, including thermodynamic values and secondary structure. (L) Dual-luciferase activity assay to determine the relationship between miR-283 and the 3’UTR of *cwo*. One-way ANOVA followed by Tukey multiple comparisons test.*** p < 0.001, ** p < 0.01, * p < 0.05.

Considering that it is miR-283 in circadian pacemaker neurons LN_v_s that plays a major regulatory role in bradyarrhythmias, we sought to investigate whether miR-283 affects heart beating by regulating the expression of biological clock genes. One of our previous studies predicted that the core rhythm genes *PAR-domain protein 1* (*Pdp1*) and *clockwork orange* (*cwo*) are potential target genes of miR-283-5p[48] and that *Pdp1* and *cwo* can be rhythmically expressed in multiple tissues[49]. We therefore further examined the expression of *Pdp1* in brain and heart, and found that *Pdp1* showed a consistent increase in transcription in senescent heart and brain, as well as increased expression in the heart of *Drosophila* with *Pdf*>mir-283^SP^ (Figure 5I), which is inconsistent with the target-binding degradation of miR-283-5p to function as a degradation mRNA.

Examination of *cwo* transcription levels revealed a significant downregulation of *cwo* in the brain and heart of aging flies and those with miR-283 knockdown in LN_v_s flies (Figure 5J), consistent with the miRNA-mRNA targeting relationship. We further predicted the potential binding targets of miR-283-5p to the 3′-untranslated region (3’ UTR) of *cwo* by the *RNAHybrid* online tool (https://bibiserv.cebitec.uni-bielefeld.de/rnahybrid) and found a potential binding site at the 489th position of the *cwo* 3’ UTR (Figure 5K). We next determined whether miR-283 could regulate *cwo* by a Dual-Luciferase Reporter Assay. The results showed that the expression of miR-283 inhibited the luciferin reporter activity of the wild-type 3’ UTR of *cwo*, but not the *cwo* mutant 3’ UTR reporter activity (Figure 5L), suggesting that miR-283 can be targeted to regulate the expression of *cwo*.

### 2.6 Exercise ameliorates bradyarrhythmias due to aging or miR-283-specific knockdown in LN_v_s

Exercise enhances cardiac health and may even partially reverse pathological cardiac remodeling in the elderly [19]. However, the underlying miRNA mechanisms by which exercise provides cardioprotection against aging are not well understood. Our study identified a role for miR-283 in LN_v_s in regulating aging-associated bradyarrhythmia and that miR-283 targeting of the core rhythm gene *cwo* may mediate this process. We next determined whether exercise improves aging heart health by modulating miR-283 expression.

First, we performed endurance exercise on *w^1118^* and miR-283 ^KO/+^ *Drosophila* for 3 weeks. Cardiac function analysis showed that exercise had a significant ameliorative effect on bradyarrhythmia due to aging in *w^1118^* flies. Specifically, exercise enhanced HR and reduced the incidence of bradycardia, and this improvement was mainly attributed to the effective shortening of DI and SI (Figure 6A-D). In addition, exercise also significantly reduced the incidence of incomplete relaxation in *w^1118^* flies (Figure 6E), suggesting that exercise exerts multifaceted benefits on cardiac function in aging *Drosophila*. However, in the miR-283^KO/+^ flies, exercise did not improve the incidence of bradycardia, although it increased HR and decreased HP by decreasing DI (Figure 6E). This result suggests that the ameliorative effect of exercise on cardiac rhythm may depend on the intact expression of miR-283.

**Figure 6.**
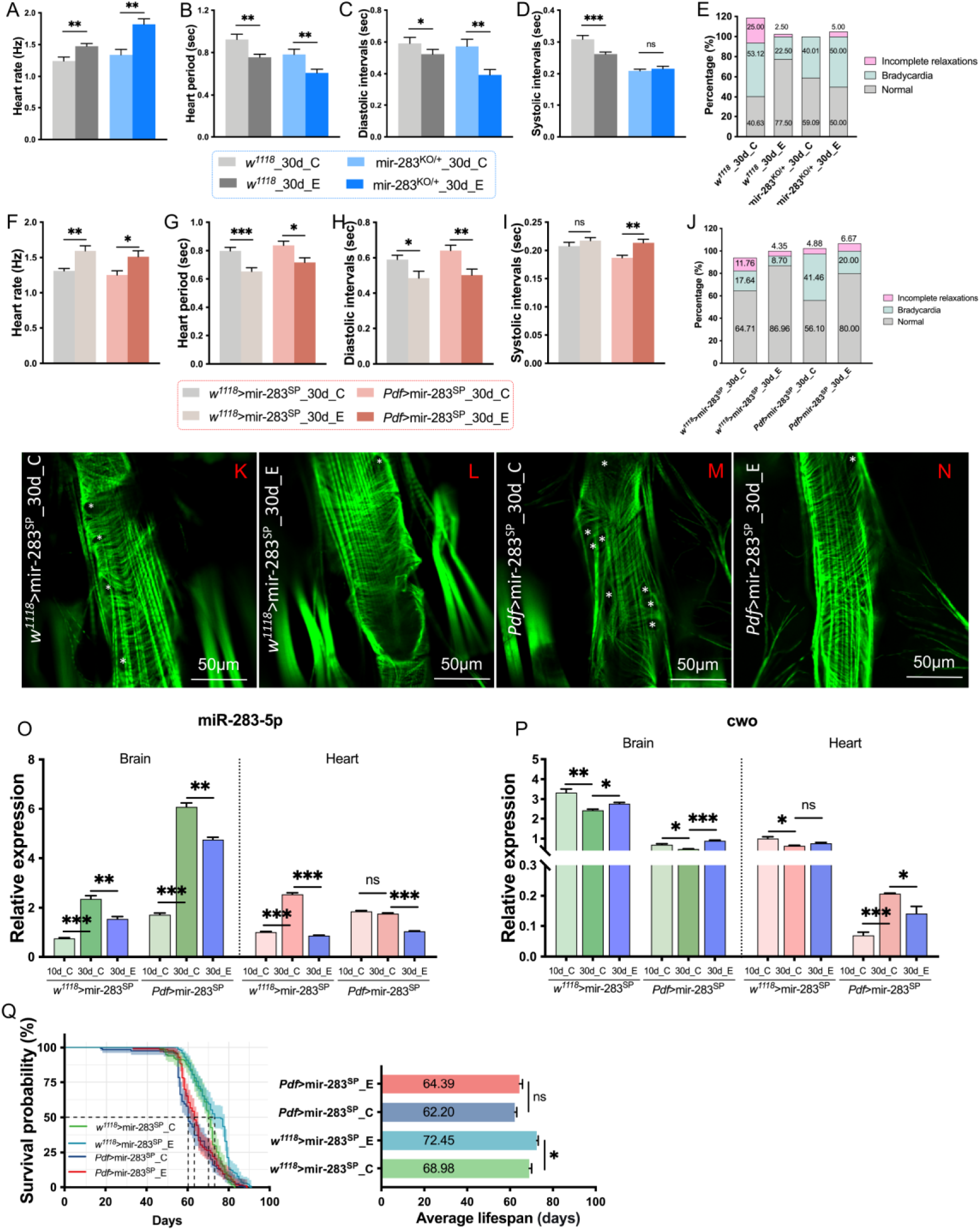
Exercise regulates miR-283 expression to improve heart health. (A-E) Heart rhythm-related indices in age-matched sedentary and exercised groups of *w^1118^* and miR-283^KO/+^ *Drosophila*. n=20-32. Student t-test. (F-J) Heart rhythm -related indices in age-matched sedentary and exercised groups of *w^1118^*>mir-283^SP^ and *Pdf*>mir-283^SP^ *Drosophila*. n=15-25. Student t-test. (K-N) Myocardial Phalloidin staining and imaging in age-matched sedentary and exercised groups of *w^1118^*>mir-283^SP^ and *Pdf*>mir-283^SP^ *Drosophila*. Significant gaps between myofibrils are indicated with “*”. n=5-8. (O-P) qPCR analysis of the expression levels of miR-283-5p and its target gene *cwo* in the brain and heart. n=20-30. Student t-test was used to analyze differences between aging (10d vs. 30d) and exercise (30d_C vs. 30d_E). (Q) Effects of exercise on the survival curves and average lifespan of *w^1118^*>mir-283^SP^ and *Pdf*>mir-283^SP^ *Drosophila*. n=120-150. Student t-test. *** p < 0.001, ** p < 0.01, * p < 0.05.

We next analyzed the effects of exercise on cardiac rhythms in *w^1118^*>mir-283^SP^ and *Pdf*>mir-283^SP^ *Drosophila*. The results showed that 3 weeks of endurance exercise was also effective in reducing bradyarrhythmia caused by aging or miR-283 knockdown in LN_v_s. The main manifestations were synchronized improvement of reduced HR, prolonged DI, and high-frequency onset bradycardia (Figure 6F-J). These improvements may be related to the fact that endurance exercise improves the morphology of contractile circumferential muscles. *Drosophila* experiencing exercise had more closely aligned ring myogenic fibers and fewer gaps than the age-matched non-exercising group (Figure 6K-N). In addition, we also observed denser non-contractile longitudinal fibers (Figure 6K-N).

Improvements in cardiac structure and arrhythmia may be due in part to the regulation of miR-283 by exercise. Exercise significantly reduced brain and myocardial miR-283 expression accumulation due to aging and miR-283 knockdown in LN_v_s (Figure 6O), and its target gene *cwo* was significantly upregulated in the brain (Figure 6P). This result suggests that the ameliorating effect of exercise on cardiac rhythm may act through nervous system miR-283-targeted regulation of *cwo*. In addition, endurance exercise significantly prolonged the mean and median lifespans of *w^1118^*>mir-283^SP^ flies (Figure 6Q). Although exercise improved lifespan in *Pdf*>mir-283^SP^ *Drosophila*, it did not reach a statistically significant difference (Figure 6Q). This provides further evidence that the overall beneficial regulatory effects of exercise on the organism may depend on the stable expression of miR-283.

In conclusion, our study suggests that 3 weeks of endurance exercise exerts a cardioprotective effect by modulating the stable expression of miR-283/*cwo* in the brain, thereby counteracting bradyarrhythmia caused by aging or miR-283 knockdown in LN_v_s.

## 3. Discussion

miRNAs play an important role in regulating deleterious changes in cardiac aging and related diseases [50], however, a limited number of role-playing miRNAs have been identified. Circulating miR-216a was found to be a circulating marker for early heart failure screening [17], and its overexpression promotes the proliferation of human cardiac fibroblasts and enhances fibrosis [51], however whether it is involved in the regulation of cardiac aging remains unknown. In this study, we systemically overexpressed or knocked out miR-283, the homolog of miR-216a in *Drosophila*, both causing age-related HR reduction and rhythm abnormalities, suggesting that the stable expression of miR-283 plays an important role in aging heart health. Further studies revealed that specific knockdown of miR-283 in LN_v_s resulted in upregulation of miR-283 throughout the brain, which in turn induced bradyarrhythmias consistent with aging. This process may be mediated by the target gene *cwo* of miR-283. In addition, inducing endurance exercise of anti gravity climbing in flies with miR-283 knockdown in LN_v_s can improve bradyarrhythmic features by decreasing brain miR-283 expression (mechanism diagram shown in Figure 7). Notably, we detected dynamic expression of miR-283, either decreasing or increasing, during different stages of cardiac aging. However, the specific knockdown of miR-283 in the heart did not affect cardiac function. This suggests that miR-283 in the heart may not be the main miRNA regulating cardiac aging or that there is compensatory regulation from the brain or other parts to maintain stable cardiac function after specific knockdown of miR-283 in the heart.

**Figure 7.**
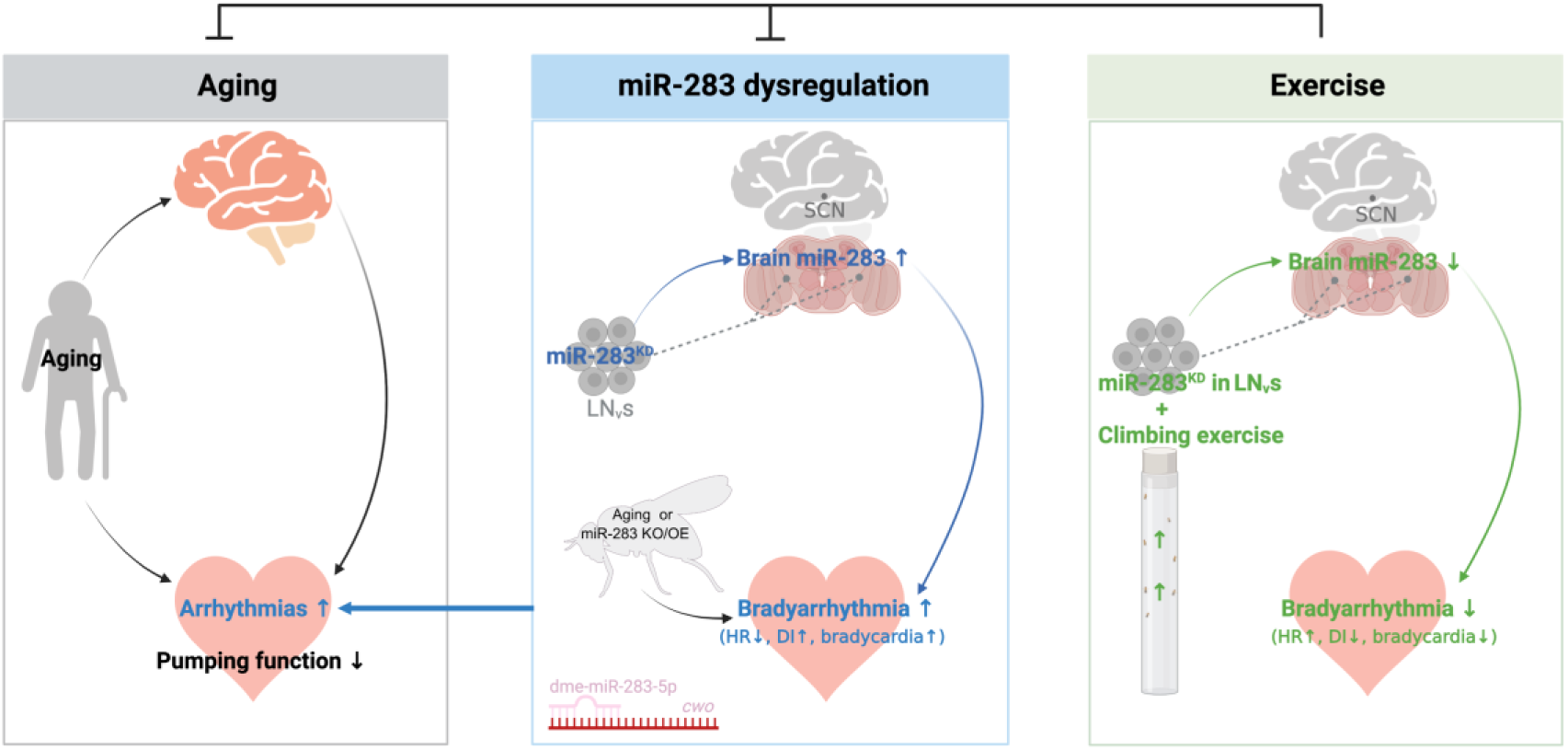
Schematic illustration of the findings obtained from this study. Based on this study, we propose that exercise ameliorates aging-associated bradyarrhythmias by modulating miR-283. First, we determined by systemic overexpression and knockout of miR-283 that abnormal expression of miR-283 can induce age-related HR reduction, prolonged DI, and increased incidence of bradycardia, which are features similar to bradyarrhythmias. Second, we specifically knocked down miR-283 in myocardium and LN_v_s, respectively, and found that miR-283 knockdown in LN_v_s leads to accumulation of miR-283 expression throughout the brain, inducing bradyarrhythmic features consistent with aging, a process that may be mediated by targeting regulation of *cwo*. Furthermore, exercise ameliorates the bradyarrhythmic features of aging or miR-283 knockdown in LN_v_s by modulating brain miR-283/*cwo* expression.

Age-related cardiovascular risks are not only closely related to the dysfunction of peripheral organ systems such as the heart and vascular system, but also closely associated with the imbalance of the autonomic nervous system[3]. Previous studies have confirmed that, neurohumoral regulation plays a critical role in cardiac remodeling to minimize changes in cardiac function [52, 53]. Thus, the autonomic nervous system plays an important role in the regulation of cardiac electrophysiology and arrhythmogenesis [54]. An in situ hybridization experiment on mir-283 showed that miR-283 is centrally expressed in the peripheral nervous system of *Drosophila* embryos, and silencing of miR-283 using the 2′Ome oligos technique induced neurological defects [55], further defining the autonomic modulatory role of miR-283. As revealed by the results of miR-283 overexpression or knockdown, the effects of miR-283 on cardiac function mainly focused on reduced HR and abnormal rhythm without affecting cardiac contractility, which predicts that the regulation of the aging heart by miR-283 may act through the autonomic nervous system. In particular, the specific knockdown of miR-283 in LN_v_s resulted in an increased range of SA-β-gal staining in the brain and heart, showing strong aging-specific changes. In addition, knockdown of miR-283 in LN_v_s resulted in disorganized myocardial myogenic fiber arrangement, and which may have contributed to more generalized bradycardia.

miRNAs regulate gene expression by inhibiting the translation of target transcripts or by inducing mRNA degradation mainly at the post-transcriptional level through sequence-specific binding [12]. In this study, we further explored the potential mechanisms by which miR-283 regulates the heart rate and rhythm. Using bioinformatics, we predicted that miR-283-5p target genes are mainly enriched in the Notch signaling pathway (*Nct*, *mam*, *Ser*, *Delta*, and *Hairless*) and could potentially regulate biological rhythm by targeting multi-tissue expression of *cwo* and *Pdp1*. Of the potential target genes tested, only *cwo* was downregulated in the heart and brain of middle-aged *w^1118^*>mir-283^SP^ and young *Pdf*>mir-283^SP^ flies, consistent with the trend for miR-283 to accumulate in senescent and LN_v_s knockdown hearts and brains and thus degrade target genes. As a transcriptional repressor, CWO can competitively bind to E-box to inhibit sustained transcriptional activation of PER-TIM by CLK-CYC [56]. Where the function of *cwo* is absent, the direct target genes of *Clk* (*per*, *tim*, *vri*, and *Pdp1*) manifest high trough values and low amplitude oscillations [57], which accounts for the significant downregulation of *cwo* and significant increase in *Pdp1* following aging and miR-283 knockdown in our results. The orthologs of the CWO in mammals are DEC1 and DEC2. In a mouse model of transverse aortic constriction-induced cardiac hypertrophy, *Dec1* deficiency protected the heart from fibrosis [58, 59]. In addition, it has been shown that DEC1 and DEC2 are involved in cardiogenesis and central nervous system development in zebrafish [60], further revealing the potential role of *cwo* in regulating cardiac function.

Exercise plays an important role in preventing cardiovascular diseases and delaying cardiovascular aging [61]. Therefore, exploring the mechanisms underlying the benign regulation of cardiac aging by exercise is necessary to reduce or reverse age-related cardiac decline. More recently, a growing number of studies have shown that miRNAs are important mediators of exercise-induced cardiac remodeling and myocardial regeneration, exerting cardioprotective effects via multiple pathways [26, 62, 63]. In our study, both aged *w^1118^* flies and *w^1118^* >mir-283^SP^ flies, as well as those with miR-283 specifically knocked down in LN_v_s, showed effective improvements in heart rate reduction and bradycardia after undergoing a three-week exercise training program. Additionally, compared to age-matched non-exercise group, the myocardial fibers of the exercised *Drosophila* were more neatly and densely arranged. Those benign improvement in cardiac function may be attributed to the lowering of accumulated miR-283 in the heart and brain by exercise, thereby relieving the inhibitory effect on *cwo*. Furthermore, we also observed a significant increase in the mean lifespan and a significant rightward shift in the survival curve in *w^1118^*>mir-283^SP^ group after exercise, suggesting that exercise produces a broad benefit in *Drosophila*.

In conclusion, our study shows that miR-283 regulates aging-associated bradyarrhythmias. In particular, miR-283 knockdown in LN_v_s, but not the myocardium, promotes the development of bradyarrhythmias and induces brain and myocardial senescence, as well as cardiac structural decline. These changes are associated with accumulation of miR-283 in the brain and heart due to miR-283 knockdown in LN_v_s, and the target gene *cwo* may mediate the regulatory effects of miR-283 on cardiac rhythm. Endurance exercise from youth effectively downregulated miR-283 accumulation in the heart and brain caused by aging or miR-283 knockdown in LN_v_s, thereby rescuing reduced heart rate and frequent bradycardia and improving myocardial structure. However, the regulatory role of miR-283 in LN_v_s on the whole brain as well as heart and even other peripheral tissues needs to be investigated in depth. In addition, we speculate that miR-283 regulation in the aging heart may act through the autonomic nervous system, but electrophysiological indicators still need to be further investigated.

## 4. Materials and Methods

### 4.1 *Drosophila* stocks and rearing conditions

*White^1118^* (*w^1118^*) was obtained from the Bloomington *Drosophila* Stock Center (BDSC#3605) as a wild-type line. MiR-283 expression regulatory strains included knockout (mir-283^KO^, BDSC#58912), overexpression (mir-283^OE^, BCF132#, y M{vas-int.Dm}ZH-2A w[*]; P{CaryP}attP40, obtained from the Core Facility of Drosophila Resource and Technology, CEMCS, CAS), and knockdown (mir-283^SP^,BDSC#61415, UAS-mCherry. mir-283.sponge). Tub-Gal4/TM3 (a gift from the Heart Development Center of Hunan Normal University) was used as a systemic driver. *Pdf*-Gal4 (BDSC#41286)[64] and *Hand*-Gal4 (BDSC#48396) driver lines are specifically expressed in ventral-lateral neurons (LN_v_s) and heart, respectively. The flies were reared on standard sugar/yeast/agar medium at a density of 30-35 flies per tube in a controlled environment at 25°C, 50% humidity, and 12 h light:12 h dark cycles (12:12h LD). The study was approved by the Ethics Committee of Hunan Normal University (No. 2024-375) and the experimental procedures were conducted in strict compliance with the ethical guidelines set by the committee. All procedures conformed to the guidelines from the Directive 2010/63/EU of the European Parliament.

### 4.2 Exercise training devices and protocols

We developed a climbing locomotion device that exploits the natural negative geotaxis of *Drosophila*. By placing the test tube (230 mm×900 mm) containing the fruit flies vertically into the card slot, the card slot rotates along its horizontal axis under the drive of the motor, and the rotation speed and angle can be set. In this study, the card slot is set for 180° vertical flip, the motor speed is 0.45 rev/s, and the reserved climbing time is 5s. Under this setting, most flies performed anti-gravity climbing movements throughout the exercise period, and a few fruit flies that failed to climb in the later period of the exercise could also actively walk up the inner wall of the test tube.

Exercise-trained *Drosophila* were subjected to a continuous 3-week exercise intervention starting from 10 days of age. Following the exercise protocol established in our previous studies[65], the flies were exercised for 2.5 hours per day for five consecutive days, followed by a two-day rest period. Data were collected at 30 days of age. The exercise sessions were conducted between ZT5 and ZT8.

### 4.3 Semi-intact *Drosophila* preparation and cardiac function

Prepare *Drosophila* semi-intact hearts using previously described methods[33]. Briefly, after anesthetizing flies with CO_2_, the head, ventral thorax, ventral abdominal cuticle, and organs were quickly removed to expose the heart. In the oxygenated artificial *Drosophila* hemolymph (ADH, 108 mM NaCl_2_, 5 mM KCl, 2 mM CaCl_2_, 8 mM MgCl_2_, 1 mM NaH_2_PO_4_, 4 mM NaHCO_3_, 15 mM 4-(2-hydroxyethyl)-1-piperazine ethane sulfonic acid, 10 mM sucrose, and 5 mM trehalose, pH 7.1), the fat around the second and third abdominal segments of the heart tube is carefully aspirated with a finely drawn glass capillarie to expose the heart edge. Then, high-speed digital movies of beating hearts were taken with a Hamamatsu EMCCD 9300 camera (Hamamatsu, Japan) at ∼124 fps for 30 s and were recorded using HCImage software (Hamamatsu, Japan). The functional cardiac parameters were assessed using the semi-automatic optical heartbeat analysis software (SOHA, kindly gifted by Ocorr and Bodmer), which can accurately detect and quantify heart rate (HR), heart period (HP), arrhythmicity index (AI, StdDev/Median of HP), systolic and diastolic intervals (SI and DI), diastolic and systolic diameters (SD and DD), and percent fractional shortening (%FS=(DD-SD/DD) ×100). Cardiac output (CO) was calculated as [π r(d)^2^ - π r(s)^2^] × HR, where r(d) is the diastolic cardiac tube radius and r(s) is the systolic cardiac tube radius[66]. In addition, a DI longer than 1 s was considered bradycardia, and a DI shorter than 0.06 s indicative of incomplete relaxations were used to estimate cardiac rhythmicity and abnormal contractions[34].

### 4.4 Fluorescence imaging

*Pdf*-Gal4 and *Hand*-Gal4 were used to drive the UAS-mCherry.mir-283.sponge *Drosophila* express mCherry fluorescence in PDF neurons, and in cardiac and paracardial cells. *Drosophila* brains [67] and semi-exposed hearts [33] were prepared according to a previously described method, and hearts were treated with hemolymph containing 10 mM EGTA after dissection to maintain a diastolic state. *Drosophila* LN_v_s and heart fluorescence expression were photographed from 3 to 5 days using a stereomicroscope (Leica DVM6, Leica Microsystems, Germany).

For F-actin staining in the heart, semi-exposed hearts were fixed with 4% paraformaldehyde at room temperature for 20 min in a relaxed state, washed with PBS three times, and then incubated with AbFluor™ 488-Phalloidin (1:500, Abbkine, BMD0082) at room temperature for 1 h, then washed 2-3 times with PBS, and mounted in an anti-fluorescence quenching agent. Confocal images were taken using a NIKON ECLIPSE TI microscope in conjunction with a NIKON C2 imaging system.

### 4.5 SA-β-gal staining and quantification

According to the manufacturer’s instructions (Senescence β-Galactosidase Staining Kit, C0602, Beyotime Bio., Shanghai, China), the dissected *Drosophila* brain and half-exposed heart were fixed in fixative solution for 30 min and washed 3 times with PBS, 5 min each. Transfer the tissue to a 1.5 ml Ep tube, add 1 ml staining working solution, and incubate overnight at 37°C. Carefully clip the brain or clip out the heart tube individually, and place it on a concave glass slide for mounting. The stained brain and heart tubes was photographed using a Leica stereomicroscope (Leica, DVM6) under the same conditions. Image J software were used to calculate the percentage of positive staining area in the whole brain or whole heart tube area by adjusting the threshold for quantitative analysis. For the analysis of the heart tube, since the accessory cardiomyocytes showed a strong β blue color, the LASSO tool in Adobe Photoshop was used to select the myocardial tissue before quantitative analysis, and the background outside the heart tube was removed.

### 4.6 Gene expression analysis

The brain, heart, and whole body of age-matched flies were collected. Total RNA was extracted using the RNA fast 200 kit (Fastagen Biotech, Shanghai, China) and cDNA was synthesized using PrimeScript™ RT reagent Kit with gDNA Eraser (TaKaRa, Japan) according to the manufacturer’s instructions. Quantitative real-time PCR (qRT-PCR) was performed on Bio-Rad CFX Connect Real-Time System with TB Green (TaKaRa, Japan) and the corresponding primer sets. The primer sequences are shown in Table S1.

### 4.7 Rapid iterative negative geotaxis (RING)

To assess the locomotion and neuromuscular coordination of *Drosophila*, negative geotaxis was performed according to the method described by Julia et al. [68, 69]. Briefly, age-appropriate flies were transferred into five cylindrical glass tubes (18 cm × 2.8 cm side by side) with sponge plugs at both ends to prevent escape and injury. During the RING process, nine shake-downs were performed, each time the fruit flies were allowed to climb for 10 s, and the complete RING was videotaped. Abandon the first three adaptive climbing times, and obtained the climbing height of the 5th seconds of 4-9 climbing videos for analysis. Since the autonomous activities of *Drosophila* show a clear rhythm with peaks of activity before and after light is turned on and off [70], we uniformly acquired RING videos during ZT11 to ZT12.

### 4.8 Survival statistics and analysis

Randomly selected 100-150 flies from each group were used for lifespan analysis. The number of dead flies was recorded at ZT14∼ZT15 each day. The average lifespan of *Drosophila* was recorded and calculated in Excel, and survival curves were plotted using the Hiplot online tool (https://hiplot.com.cn/home/index.html).

### 4.9 Bioinformatics analysis

The DIANA microT-CDS online tool (http://diana.imis.athena-innovation.gr/DianaTools/) was used to predict target genes for dme-miR-283-5p. The DAVID Bioinformatics Resources (https://david.ncifcrf.gov/) was utilized to perform pathway enrichment analysis of the predicted target genes. Use the online resource RNAhybrid (https://bibiserv.cebitec.uni-bielefeld.de/rnahybrid) to predict the binding targets of dme-miR-283-5p and corresponding mRNA 3’UTR, and generate a secondary structure.

### 4.10 Cell culture, transfection, and luciferase assay

293T cells (human embryonic kidney cells) were cultured in DMEM (BasalMedia, Shanghai, China) containing 10% FBS and 1% P/S under conditions of 37°C and 5% CO_2_. The reporter genes for the wild-type 3’UTR of *cwo* and the mutant 3’UTR with predicted binding sites were cloned into the multiple cloning sites of the psiCheck-2 vector (1643-1674). Both the plasmids and miR-283 mimics were synthesized by Tsingke Biotech Co. (Beijing, China). The transfection process of plasmids and miRNA mimics was mediated using Lipo8000^TM^ transfection reagent (Beyotime Biotechnology Co., Shanghai, China) in a 96-well plate. The transfection amount of plasmids was 100 ng / well, and the transfection amount of miRNA was 5 pmol / well. Luciferase activity was measured according to the instructions of the Dual-Luciferase^®^Reporter (DLR™) Assay System (Promega, UAS) and normalized to the Firefly luciferase data.

### 4.11 Statistical analysis

Data were processed and statistical analysis using SPSS 22.0. In the SOHA analysis, data were normalized using the Z-score normalization method prior to data analysis, and abnormal data with absolute values ≥ 3 were removed. Subsequently, data that conformed to a normal distribution and variance homogeneity were subjected to an independent samples t-test or one-way ANOVA with Tukey’s multiple comparison test. Two-way ANOVA was used to detect the interaction effects of miR-283 knockout and aging, as well as the specific knockdown of miR-283 in LN_v_s and aging, multiple comparisons were performed using Tukey’s test. GraphPad Prism was used for graph generatio and is displayed as mean ± SEM. Statistical significance was set at at *p* < 0.05.

## Author Contribution

Qiufang Li: Conceptualization; methodology; data curation; formal analysis; visualization; writing—original draft. Lan Zheng: Conceptualization; resources; supervision; funding acquisition; project administration; writing—review and editing. Xu Ping: Data curation; formal analysis; Zhengwen Yu: Data curation; formal analysis. Qin Yi: Data curation; formal analysis. Chao Tang: Data curation. Xiaoya Wang: Data curation.

## Acknowledgements

We are grateful to the Center for Heart Development of Hunan Normal University for providing Tub-Gal4/TM3 flies.

